# Optogenetic stimulation of medial septal glutamatergic neurons modulates theta-gamma coupling in the hippocampus

**DOI:** 10.1101/2022.03.15.484394

**Authors:** Elena Dmitrieva, Anton Malkov

## Abstract

The oscillatory activity of the hippocampus and parahippocampal structures encodes information during cognition. Multiple researchers emphasize hippocampal cross-frequency theta-gamma coupling (TGC) as a basic mechanism for information processing, retrieval, and consolidation of long-term and working memory. While the role of entorhinal afferents in the modulation of hippocampal TGC is widely accepted, the influence of other main input to the hippocampus, from the medial septal area (MSA, the pacemaker of the hippocampal theta rhythm) is poorly understood. This area innervates multiple targets in the hippocampus, so information about the participation of MSA in cross-frequency interactions might be critical for an understanding of neural coding.

Optogenetics allows us to explore how different neuronal populations of septohippocampal circuits control neuronal oscillations in vivo. It was shown that optogenetic activation of glutamatergic projections to the hippocampus with theta-frequency stimuli paces the hippocampal theta rhythm. Here we investigated the influence of phasic activation of MSA glutamatergic neurons expressing channelrhodopsin II on theta-gamma coupling in the hippocampus. Transgenic mice B6.Cg-Tg (Thy1-COP4/EYFP)18Gfng/J were used in this experiment. Expression of ChRII in MSA was verified immunohistochemically and the optic fibers just above MSA and the recording electrodes in the CA1 hippocampal field were implanted. During the experiment, local field potentials of MSA and hippocampus of freely behaving mice were modulated by 470nm light flashes with theta frequency (2-10) Hz.

It was shown that both the power and the strength of modulation of gamma rhythm nested on hippocampal theta waves depend on the frequency of stimulation. The modulation of the amplitude of slow gamma rhythm (30-50 Hz) prevailed over modulation of fast gamma (55-100 Hz) during flash trains and the observed effects were specific for theta stimulation of MSA. We discuss the possibility that phasic depolarization of septal glutamatergic neurons controls theta-gamma coupling in the hippocampus and plays a role in memory retrieval and consolidation.

## Introduction

The brain uses neuronal oscillations to organize information processing, coding and retention. Different neuronal populations and brain structures produce rhythmic fluctuations of the neuronal membrane potential, which serve as a temporal window for communication. The hypothesis “communication through coherence” (CTC) (Fries, 2005) postulates that only coherently oscillating (at the same frequency) neuronal groups can interact effectively, because their communication windows for input and for output are open at the same time. It has been shown that phase coherence reflects various cognitive processes in humans (Canolty et al., 2006; Axmacher et al., 2010) and animals (Montgomery and Buzsáki, 2007; Tort et al., 2008, 2009; Wulff et al., 2009; Canolty et al., 2010). Moreover, in these and many other studies rhythms of different frequencies coexisted and were often synchronized to each other or nested into each other. The cross-frequency coupling may represent a mechanism for the regulation of communications between different spatiotemporal scales (Palva et al., 2005; Holz et al., 2010). The phase coupling between theta and gamma oscillations, namely, the modulation of the gamma amplitude by theta phase, is the most studied phenomena of phase coherence (Buzsáki et al, 2003; Bragin et al., 1995; Mormann et al., 2005; Canolty et al., 2006; Sirota et al., 2008; Tort et al., 2008; Schomburg et al., 2014).

Theta-rhythm (4-12Hz), the principal oscillatory pattern generated by the hippocampus, plays a fundamental role in cognitive processes such as learning, spatial navigation, memory consolidation, reaction to novelty, selective attention and many more. A detailed analysis of the commonality of all these functions suggests that the theta rhythm serves as a temporal filter that provides admittance to the output of hippocampal neurons only those signals that come in certain phase relation to the theta wave (Vinogradova, 2001). All neuronal populations of the hippocampal formation, to a greater or lesser extent, show binding of their activity to the phase of the theta-rhythm, in other words, they generate action potentials predominantly in a certain phase of the theta cycle (Klausberger et al., 2003; Mizuseki et al., 2009; Somogyi et al.,2014), thus forming the nested high-frequency oscillations. A bulk of evidence suggests that successful memory performance requires coupling of 30–100 Hz gamma rhythms to particular phases of the theta cycle (Tort et al., 2009; Igarashi et al., 2014; Takahashi et al., 2014; Colgin et al., 2009). It is assumed that the mechanisms of organization of slow (~25-50 Hz) and fast (~55-100 Hz) gamma-oscillations are different since they are generated by different networks involving some particular classes of GABAergic interneurons (for review, see Colgin, 2016; Mably and Colgin, 2018). Gamma patterns segregate around theta phase: slow gamma dominated CA1 local field potentials (LFPs) on the descending phase of CA1 theta waves during navigation, and fast gamma was bound to the theta peak (Hasselmo et al., 2002; Shomburg et al., 2014). These signals corresponded to CA3 and entorhinal input, respectively. Long-term potentiation in CA1 is most easily induced at the ascending phase of theta when EC input is maximal. This indicates that the theta phase when EC inputs preferentially arrive may coincide with the time when memory encoding occurs optimally and raises the possibility that the EC related CA1 fast gamma facilitates memory encoding. Retrieval of information is thought to occur at a different theta phase than memory encoding, during which time CA3 input to CA1 is maximal and incoming signals from EC are suppressed (Colgin et al.,2009).

While the role of entorhinal afferents in the modulation of hippocampal TGC is widely accepted, the influence of other main input to the hippocampus, from the medial septal area (MSA) is poorly understood.

Hippocampus is considered as the main and the only generator of the theta rhythm; however, its activity shows a strong dependence on the connection with MSA. Lesion of MSA or its projections leads to the disruption of the theta activity in the hippocampus (Gray, 1971; Mitchel et al., 1982). A less structurally organized MSA, whose neurons fire in bursts at the theta frequency plays a role of a pacemaker of the hippocampal theta rhythm.

MSA contains three main types of neurons: GABAergic, cholinergic, and glutamatergic neurons (Manseau et al, 2005; Sotty et al., 2003). One population of GABAergic neurons expressing calretinin innervates other cells within the MSA only (Kiss et al., 1997). The second group of neurons projects to the hippocampus and expresses parvalbumin (Freund, 1989). There is a consensus that it is projections from parvalbumin-containing GABAergic neurons terminating on hippocampal interneurons that are involved in synchronization of the septo-hippocampal network at theta frequency (Hangya et al., 2009; Varga et al., 2008). Silencing of MSA GABAergic neurons suppresses theta-gamma coupling and theta phase coherence (Bandarabadi et al., 2017).

MSA forms the main cholinergic input to the hippocampus. The role of cholinergic neurons in the generation of theta rhythm and cross-frequency coherence in the hippocampus was demonstrated using an optogenetic approach that allows selective stimulation of various neurochemical populations of MSA. Medial septal cholinergic activation can both enhance theta rhythm and suppress peri-theta frequency bands, allowing theta oscillations to dominate (Vandecasteele et al., 2014). In addition, activation of cholinergic MSA neurons led to a synergistic suppression of the activity of CA3 pyramids due to direct metabotropic activation of hippocampal interneurons and, at the same time, an indirect effect due to excitation of septal parvalbumin projections. As a result the suppression of the slow gamma input by CA3 inhibition created conditions for fast gamma coupling and information encoding in CA1 (Dannenberg, 2015).

Glutamatergic MSA neurons synaptically drive hippocampal principal cells (Huh et al., 2010). Robinson and coauthors (2016) showed that while hippocampal oscillations strictly followed to the frequency of optogenetic stimulation of MSA glutamatergic neurons in theta range, activation of septo-hippocampal glutamatergic terminal had no effect on theta rhythm. The role of septal glutamatergic neurons in cross-frequency coupling has not been previously studied.

Here we investigate how phasic activation of MSA glutamatergic neurons in different frequencies influences theta-gamma coupling in the hippocampus.

## Materials and methods

### Animals

B6.Cg-Tg(Thy1-COP4/EYFP)18Gfng/J mice(n=7, 30-35g) were used for experiments. The animals were acclimatized for 1 week prior to the start of the experiments. Mice were housed in cages with free access to clean drinking water, 3-4 animals each, under a standard 12-hour light/dark cycle at a temperature of 22 ± 2°C and at a humidity of 45% ± 5%. The study was conducted in accordance with the ethical principles formulated in the Helsinki Declaration on the care and use of laboratory animals and the Regulations of the European Parliament (86/609/EC).

### Surgery

A series of neurosurgical operations were performed on the implantation of recording electrodes - nichrome wire 15 microns, as well as the implantation of a stimulating optical fiber. The operation was performed under general combined anesthesia using tiletamine zolazepam (Zoletil 35 mg/kg) as the main anesthetic and xylazine (25 mg/kg) as a muscle relaxant. After the onset of the deep stage of sedation, the animals were fixed in a stereotaxic apparatus. After scalping, the cranial bone was dried (70%, 96% alcohol), treated with the hemostatic drug. Electrodes for recording field activity were implanted unilaterally according to pre-calculated coordinates in hippocampal field CA1 (AB = −2.5; L = 2; H = 1.8), hippocampal field CA3 (AB = −2.8; L = 3.2; H = 3.8), and medial septum. (AB = + 0.8; L = 1.2; H = 4; 15°). The optical fiber was implanted into the medial septum (AB = + 0.8; L = 1.2; H = 4; 10°) or to the entorhinal cortex (AB = −3.7; L = 4.5; H = 4.8). Electrode and fiber placement was verified histologically after the experiments. The reference electrode was screwed into the occipital bone above the cerebellum. The entire complex was fixed on the head with dental acrylic. After the operation, the wound surface was treated with a 5% alcohol iodine solution.

### Optical stimulation in vivo

After a recovery period of three days, a series of optical stimulations were performed on. The setup for optical stimulation (ThorLab, USA) consists of a light source with a wavelength of 470 nm, which is connected to a controller. Optic fibers with a diameter of 2.5 mm are connected by a rotating contact and attached to the cannula before the experiment. Before the start of the test, the animals adapted to the experimental environment for 5-10 minutes. The animals moved freely in the chamber. After the allotted time, burst stimulation was applied to the fiber with pulses 10-20 ms long in the range 2-12 Hz, as well as stimulation 30 Hz for one minute. The interval between stimulations was at least 10 minutes, 2-4 stimulations per experimental session. Field potentials were recorded using a custom made amplifier and ADC Instrutech LIH8+8 (sampling frequency 5 kHz). Data recording was carried out using the WinEDR 3.7 software.

### Histology

After the experimental series, the animal was decapitated. For immunochemical analysis of the brain, 30 μm sections prepared on a cryotome were placed in a Petri dish with 1–2 ml of 0.1 M phosphate buffer with 0.3% Triton X-100 detergent. After washing, the sections were placed in a blocking solution with 1% BSA for 2 hours. After incubation, the sections were washed with a buffer and antibodies to VGluT2 + Alexa Fluor 647 (Novus Biologicals) were added to them for 2 hours. The sections were again washed with a buffer and placed on a glass slide. The level of fluorescence was checked under a Nikon E200 microscope, with fluorescent filters Ex/Em =470/530nm for visualization of YFP associated with channelorhodopsin II, and Ex/Em =620/650nm for Alexa Fluor 647.

### Data analysis

Data analysis was carried out using the IGOR PRO software. For frequency analysis the Fast Fourier transform was used. Wavelet transform was used to determine the current phase and amplitude of the oscillations. Next, high-amplitude signal components were identified in the 30– 100 Hz band with a step of 1 Hz, exceeding the standard deviation by 3 or more times. The integrated amplitude of the selected components was normalized for each frequency and used to construct cross-frequency and phase-amplitude diagrams. The modulation index was calculated equal to the Kullback-Leibler distance between the distribution of the amplitude of the gamma rhythm in the theta phase and the distribution of the unmodulated signal.

The significance of changes during stimulations was assessed using the Mann-Whitney U-test for independent samples, as well as using ANOVA analysis of variance for repeated measurements. The results of the studies are presented as a result ± SEM, where SEM is the standard error of the mean.

## Results

### ChR-2-YFP is expressed in glutamatergic MSA neurons

In our experiments, we used B6.Cg-Tg(Thy1-COP4/EYFP)18Gfng/J mice. Thy1 is a neuronal marker and is most commonly used to study excitatory pyramidal cells in the hippocampus and cortex. Although many studies (Dobbins et al. 2018; Feng et al. 2000) indicate that Thy1 is a marker of glutamatergic neurons, co-expression of ChR-2-YFP and VGLUT2 in MSA has not been previously confirmed. After MSA immunohistochemical staining, we were able to demonstrate co-expression of ChR-2-YFP and VGLUT2, in other words, all YFP-expressing MSA neurons were glutamatergic (Fig.1A).

**Fig.1.**
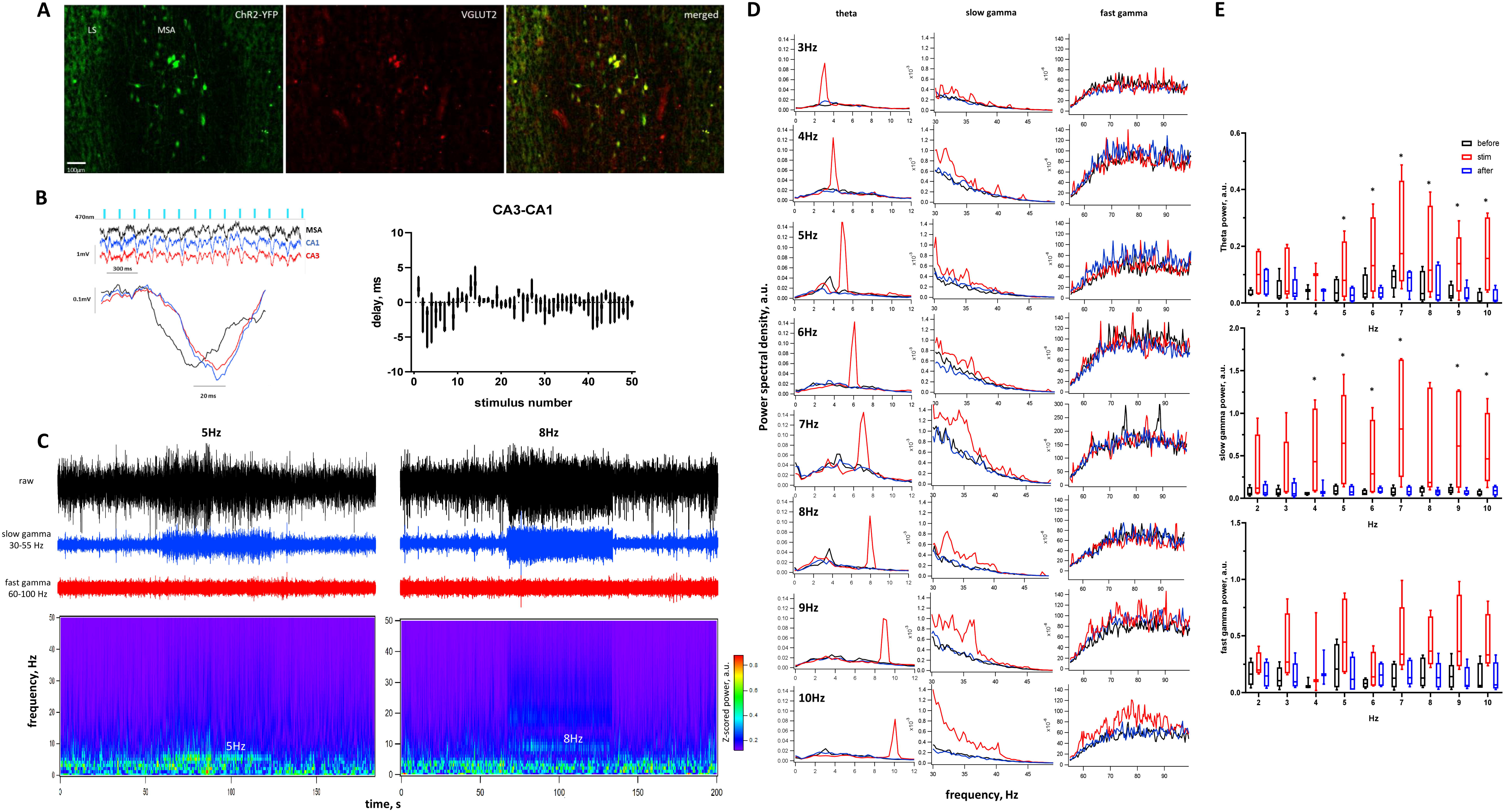
A) Immunofluorescence images from brain slices show co-expression of ChR2-YFP (green) and VGLUT2 in the medial septal area of a transgenic mouse. B)Left-LFP traces and single responses to light stimulation in MSA, CA1 and CA3, note time delay between depolarization of MSA and hippocampus. Right – Min-Max boxplots showing time delay between CA3 and CA1 to first 50 light stimuli. C. Raw (black) and filtered (blue - slow gamma band, red – fast gamma) records of hippocampal LFP during optical stimulation of MSA at 5 and 8 Hz. D) Power spectral density (PSD) graphs of LFPs from CA1 before(black), during(red) and after (blue) optical stimulation of different frequencies. E) Min-Max boxplots showing integrated (PSD) depending on the frequency of stimulation for theta (top), slow gamma (middle) and fast gamma (bottom) bands.

### Optical stimulation drives theta activity

We first determined how optical stimulation affects the field potentials of the MSA and hippocampal CA1 and CA3 fields. Both structures responded with pronounced depolarization to a single flash of light (Fig.1B). Besides, a pronounced synaptic delay was observed between the MSA and hippocampal fields. Depolarization of septal neurons was ahead of the theta wave in the hippocampus by 18±6 ms. The depolarization responses of the CA3 and CA1 fields were mostly synchronous. Analysis of the delay between fields CA3 and CA1 during the stimuli train showed that the value is maximal (CA3 ahead by 2.8±1.8 ms) at the beginning of stimulation, and gradually decreases to values close to zero (Fig.1B).

As previously discovered by Robinson et al. (2016), we have shown that the rhythmic activity of the hippocampus and MSO is related to the frequency of optical stimulation in the theta range. When stimulated in the range from 2 to 10 Hz for 1 min with a duration of light flashes of 10–20 ms, a clear correlation was observed between the frequency of rhythmic activity of the hippocampus and the frequency of stimulation. The effect disappeared immediately after the stimulation was turned off. Fig.1C shows natural recordings of hippocampal local field potentials before, during and after optical stimulation at 5 and 8 Hz MSO: pronounced responses to stimulation in the hippocampus are shown.

The power of induced theta rhythm depended on the frequency of stimulation. Theta-band power was significantly higher than control values during stimulation in the range 5-10 Hz with maximum at 7 Hz (275±96%, p<0.05, Fig.1 D,E). Slow gamma oscillations also increased significantly (Fig.1C,D). Power changes were most pronounced at 5Hz (724±262%, p<0.01) and 7Hz (915±390%, p<0.01, Fig.1E). Fast gamma rhythm was higher compared to control, but we were unable to detect significant changes in the power during stimulation. Peak frequencies in the gamma range did not depend on the stimulation frequency, so we excluded the effect of stimulation harmonics on the observed effects. No behavioral changes such as freezing, grooming, or hyperactivity were detected during stimulation.

### Theta-gamma coupling during stimulation of glutamatergic septal neurons

In our experiments, slow and fast gamma rhythms in the CA1 field of hippocampus were nested in the ascending and descending phases of the theta cycle, respectively, which agrees with the earlier data (Buzsáki et al., 1983; Bragin et al., 1995; Belluscio et al., 2012).

Optogenetic activation of glutamatergic MSA neurons significantly transformed the TGC pattern in the hippocampus. Stimulation at frequencies in the theta range led to the dominance of the slow gamma coupling in all animals (Fig. 2). The maximum values of the modulation index (MI) of slow gamma oscillations were observed at a stimulation frequency of 5 Hz. An analysis of cross-frequency modulation showed that the slow gamma is modulated precisely by the theta stimulation frequency (Fig. 2C). Also, in all experiments, a slow gamma rhythm was modulated over the entire range of 30-55 Hz, regardless of the stimulation frequency.

**Fig.2.**
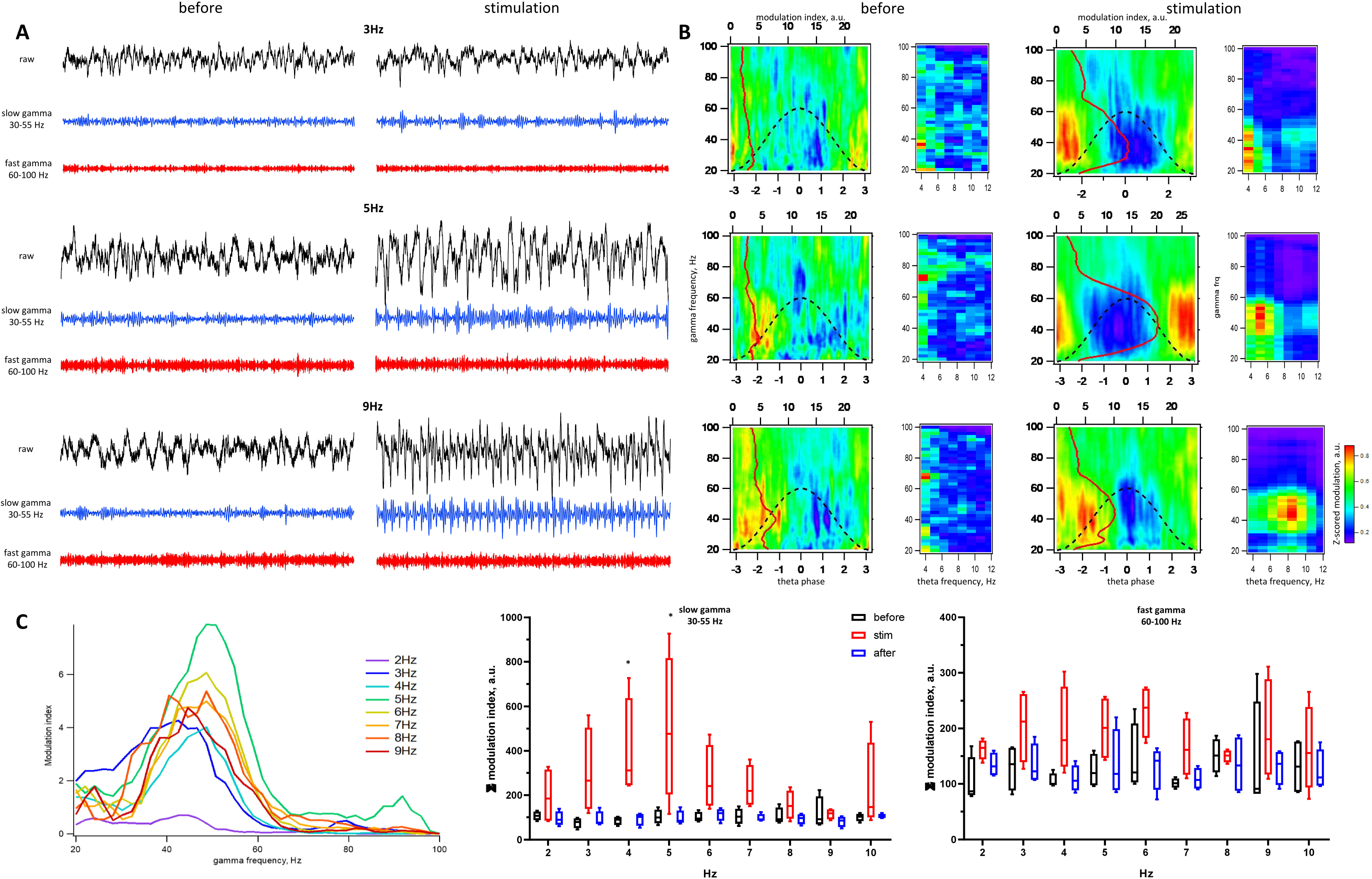
A) Raw (black) and filtered (blue - slow gamma band, red – fast gamma) records of hippocampal LFP before and during optical stimulation with the frequency 3,5 and 9 Hz. B) Phase-amplitude (first and third column) and cross-frequency (second and fourth column) modulation diagrams of traces presented on (A). Z-scored modulation index shown by color. Red line (and top axis) shows the modulation index profile for gamma frequencies. C) Left-Mean modulation index profiles of LFPs during stimulation of different frequencies. Middle and Right - Min-Max boxplots showing integrated modulation index before(black), during(red) and after (blue) optical stimulation of different frequencies of slow and fast gamma bands.

Unlike slow gamma, fast gamma did not show any significant changes during stimulation (Figure 2C).

To verify the specificity of the observed effects, experiments were performed with MSA stimulation at a frequency of 30 Hz, as well as with optical theta activation of the medial entorhinal cortex. In both cases, there was no effect on slow gamma TGC, as was the case with MSA theta stimulation. (Fig. 3)

**Fig.3).**
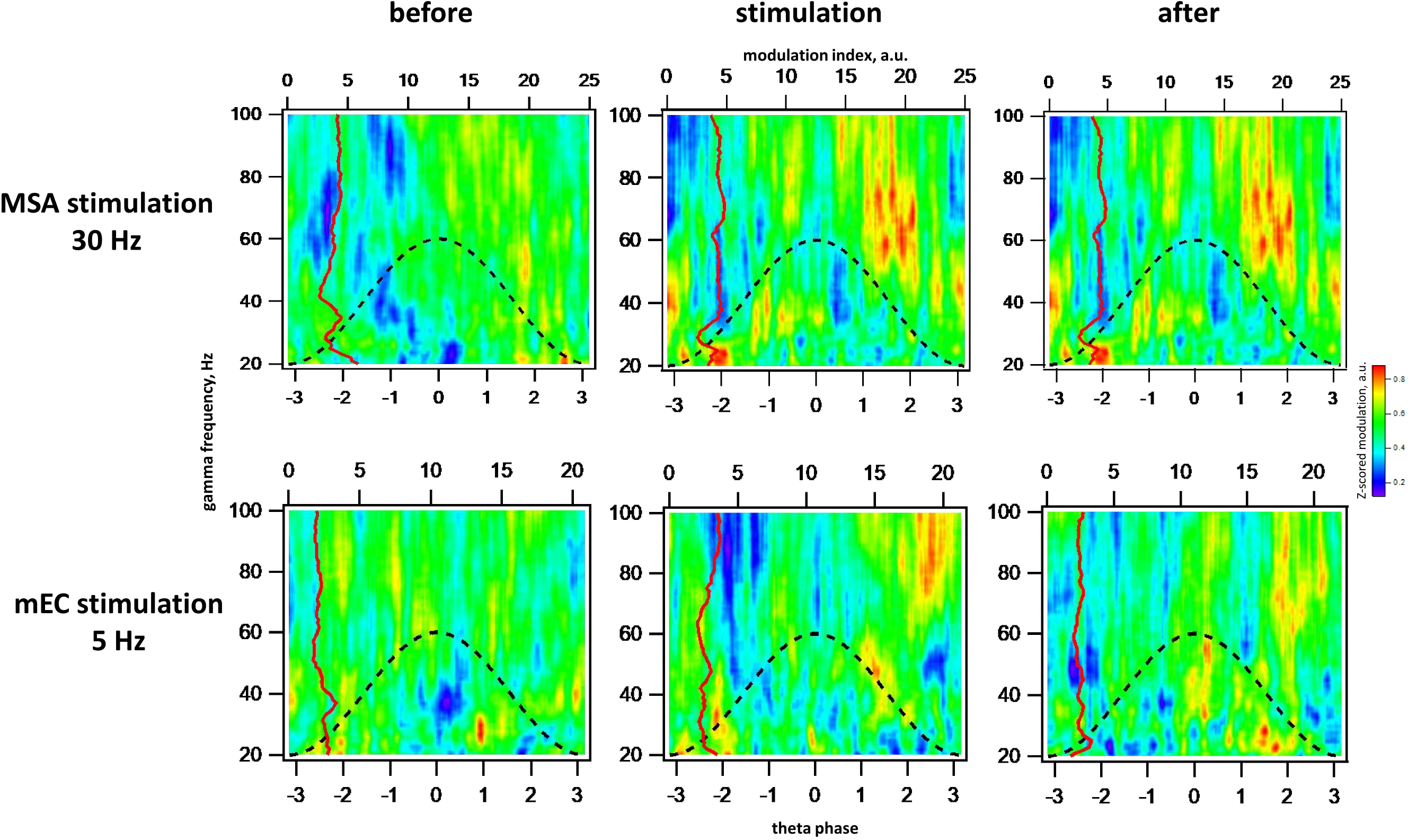
Phase-amplitude modulation diagrams for 30 Hz MSA (top) and 5 Hz mEC (bottom) stimulations. Note the absence of effect on slow gamma coupling.

## Discussion

Medial septal area, the pacemaker of hippocampal theta rhythm, forms a main subcortical input to the hippocampus. Neuronal populations of MSA are known to play a critical role in control of hippocampal activity and excitability. However, it is still unknown how these populations interact with each other and what is their role in information processing in the hippocampus. Here we investigated how septal glutamatergic neurons are involved in generating hippocampal theta activity and phase-amplitude coupling of theta and gamma rhythms.

We used an optogenetic approach to rhythmically depolarize glutamatergic MSA neurons and assessed the basic parameters of hippocampal oscillations. As in earlier work by Robinson and co-authors (2016). We have demonstrated that such stimulation leads to a tight tuning of hippocampal activity in the theta band. The authors ruled out a direct effect of glutamatergic projections and suggested that this tuning is associated with modulation of the activity of GABAergic neurons within MSA. Relatively long synaptic delay between septal and hippocampal depolarization in our experiments does not exclude the presence of polysynaptic transmission. Nevertheless, it should be noted, that at the very first field responses of the hippocampus, the CA3 field depolarized earlier, indicating that direct activation of the pyramidal neurons takes place.

Unlike in the mentioned study, we were able to demonstrate the dependence of the power of the evoked theta rhythm on the stimulation frequency. The high sensitivity of the hippocampus to a stimulation frequency of 7 Hz can be explained by intrinsic mechanisms of theta rhythm generation. Interneurons in CA1 are characterized by the presence of a slowly inactivating potassium current that occurs during hyperpolarization. This current does not depend on synaptic inputs and causes rhythmic fluctuations in the membrane potential in the range of 5–7 Hz (Chapman and Lacaille, 1999). Also, there is evidence that a prolonged (2 sec) incoming depolarization current activates high-threshold calcium channels, which causes oscillations of the membrane of pyramidal neurons at a frequency of 7 Hz and below (Garcia-Mufioz, et al., 1993). Thus, the existence of intrahippocampal mechanisms for generating theta oscillations suggests that the optimal modulation of theta activity is excitation or inhibition at frequencies of 5-7 Hz.

It is believed, that slow gamma oscillations coupled to theta wave promote the retrieval of stored memories (Tort et al., 2009; Shirvalkar et al., 2010; Igarashi et al., 2014) while the fast gamma rhythm facilitates the encoding of current signals (Montgomery and Buzsaki, 2007; Colgin et al., 2009; Newman et al., 2013). Analysis of cross-frequency modulation in our experiments revealed a sizable increase in theta-slow gamma coupling. A necessary condition for the occurrence of slow gamma coupling is the activation of intrahippocampal connections between CA3 and CA1 through Schaffer collaterals (Hasselmo, 2002; Schomburg et al.,2014; Colgin,2015). Recently it has been shown in slices that direct optogenetic activation of CA3 principal cells induces a single theta wave with nested slow gamma oscillations, while other stimulation sites respond by higher frequencies (Butler et al., 2018). Moreover, as mentioned above inhibition of CA3 activity by activation of cholinergic MSA neurons suppresses slow gamma and creates conditions for fast gamma coupling in CA1 (Dannenberg, 2015). MSA glutamatergic neurons are known to innervate a portion of both interneurons and pyramidal neurons in the hippocampus (Huh et al., 2010; Sun et al., 2014). Thus, we propose that rhythmic activation of glutamatergic septal projections to the CA3 pyramids may induce the theta-slow gamma coupling in the hippocampus. Antagonism of septal glutamatergic and cholinergic neurons may be an important mechanism for involvement of MSA in information processing and triggering between information encoding and retrieval in the hippocampus.

A detailed understanding of the contribution of MSA to cross-frequency modulation in the hippocampus requires further studies to assess the involvement of GABAergic neurons and to describe how different neuronal populations interact during information processing.

## Acknowledgments

The work was supported by the Russian Science Foundation (project No. 22-25-00475).

